# Effects of base temperature, immersion medium, and EM grid material on devitrification thresholds in cryogenic optical super-resolution microscopy

**DOI:** 10.1101/2025.03.28.645948

**Authors:** Soheil Mojiri, Joseph M. Dobbs, Niko Faul, Thomas P. Burg, Julia Mahamid, Jonas Ries

**Author notes:** Corresponding authors. Email addresses (Julia Mahamid), (Jonas Ries).

## Abstract

Cryogenic correlative light and electron microscopy (cryo-CLEM) is an imaging strategy that integrates specific molecular labeling and molecular resolution structural information. However, there is a resolution gap of more than two orders of magnitude between diffraction-limited fluorescence microscopy and electron microscopy (EM). Single-molecule localization microscopy (SMLM) performed at cryogenic temperatures promises to bridge this resolution gap. Nevertheless, the high excitation laser powers required for SMLM risk the devitrification of frozen biological samples, leading to perturbation of their native-like state. Here, we investigate how base cooling temperature, immersion medium, and EM grid support materials influence sample devitrification. Using finite element simulations and experimental validation, we show that a cryo-immersion medium enhances heat dissipation for carbon supports, while metallic supports in a cold nitrogen gas medium tolerate higher laser intensities due to lower base temperatures. Gold supports illuminated at 640 nm exhibit markedly high laser thresholds, similar to silver-coated grids. Additionally, metallic supports maintain efficient heat dissipation in vacuum-based cryostats. Our findings provide quantitative insights that aid in optimization of cryo-SMLM setups for improved cryo-CLEM imaging.

## 1. Introduction

Cryogenic electron microscopy (cryo-EM) is a valuable tool in structural and cell biology [1–3]. In particular, cryo-electron tomography (cryo-ET) allows high-resolution, three-dimensional visualization of the complex organization and interactions of macromolecules and organelles within intact cells. However, identification of specific macromolecular complexes in cryo-ET data remains a challenging task. Only complexes larger than approximately 300 kDa produce sufficient signals in the cryo-ET data for direct identification based on their structural signatures [4].

Under cryogenic conditions, the combination of fluorescence and electron microscopy, known as cryo correlative light and electron microscopy (cryo-CLEM), merges the benefits of high-content, high-resolution electron microscopy images with the specificity of a fluorescent tag on a protein target [5, 6]. However, a resolution gap of more than two orders of magnitude exists between diffraction-limited cryo light microscopy (450*−*600 nm in lateral and 2 µm in axial dimensions [7, 8]) and electron microscopy, which in many cases prevents the unambiguous and precise localization of a specific molecule. Super-resolution cryo-CLEM (SR-cryo-CLEM) overcomes this challenge by enhancing the spatial resolution of fluorescence imaging.

The majority of existing SR-cryo-CLEM implementations utilize single-molecule localization microscopy (SMLM) techniques, in particular, photo-activated localization microscopy (PALM) [9–15]. In PALM, a long movie of blinking fluorophores is acquired where only a small subset of fluorescent molecules are switched from a non-emitting to a photon-emitting state per image frame using photo-activation to enable single-molecule localization. The localization precisions reported under cryogenic conditions using photoactivable fluorescent proteins, ranging from about 10 to 30 nm [10–13, 15, 16], are mainly limited by the number of collected photons and the mechanical stability of the microscopes.

The performance of cryo-SMLM involves several primary yet interconnected challenges. First, exposing biological samples vitrified on conventional EM grids to high excitation laser intensities, similar to those used in room temperature SMLM (1-500 kW/cm^2^ [17]), raises the ice temperature above 138 K, leading to devitrification and ultimately sublimation. SR-cryo-CLEM techniques that use lower excitation intensities, such as super-resolution optical fluctuation imaging (SOFI) [18], structured illumination microscopy (SIM) [8, 19], and image scanning microscopy (ISM) [20], generally enhance the resolution only two to three times compared to conventional diffraction-limited imaging. Second, to collect sufficient photons, the required low-intensity excitation intensities lead to long image acquisition times. This is exacerbated by the fact that fluorophores tend to remain in the emitting state longer at cryogenic temperatures [13], where photobleaching rates can be up to two orders of magnitude lower compared to room temperature [21–23]. This amplifies the impact of mechanical instabilities and increases the risk of ice contamination within cryo-stages [24, 25].

Therefore, accurate characterization of the parameters that influence devitrification is of paramount importance to optimize cryo-SMLM imaging for cryo-CLEM applications. Dahlberg et al. demonstrated that a vitrified sample can be safely illuminated with up to 1 kW/cm^2^ by coating conventional EM grids with a thin film of silver, highlighting that the light absorption and heat conductivity properties of the support film have a substantial impact on the devitrification threshold [26]. In addition to confirming the significant role of the support film material, Last et al. characterized the effect of the grid material, grid geometry, film geometry, film thickness, and ice thickness on the devitrification threshold [27]. The above-mentioned studies exploit open (non-vacuum) LN2-cooled cryo-stages operated at 77 K base temperature and utilize cold N2 gas as the sample immersion medium.

Devitrification thresholds in closed (vacuum) cryo-stages operated at different base temperatures (77 K or 8 K [28, 29], 89 K [30]) have not been investigated in detail. Similarly, little information is available for open (non-vacuum) cryo-stages that utilize different liquid immersion media (liquid propane [22], Ethoxynonafluorobutane (HFE-7200) [31, 32], 70:30% methanol:1-propanol [33]), and correspondingly different base temperatures of 100 K, 133 K, and 128 K. Notably, depending on stage design, the actual base temperature in a variety of LN2-cooled stages is usually higher than 77 K (e.g. 113 K [10], 98 K [19], 93 K [11], and 80 K [9]). This may represent one source of discrepancy in the reported laser intensity thresholds in different studies.

In this work, we quantified how the cryo-stage base temperature and immersion medium affect laser intensity-based devitrification thresholds using heat transfer simulations. We identified the highest safe laser intensities for conventional and custom-coated grids and validated our findings by illuminating vitrified *Escherichia coli* (*E coli*) cells. Experiments were conducted using a cryo-immersion (HFE-7200) microscope at 133 K, a LN2-cooled open cryostat at 95 K, and a LN2-cooled closed (vacuum) cryostat at 77 K. Finally, we extended our simulations to vacuum-based cryo-stages operated at ultra-low liquid helium (LHe) temperatures (8 K) to explore the same parameters.

## 2. Results

Transmission electron microscopy (TEM) grids are typically thin, flat, and circular, with a standard 3 mm diameter. In cryo-EM applications, the grid mesh is commonly made from copper or gold. The mesh size determines the size of each square (e.g. 65 µm for 300 mesh grids, 90 µm for 200 mesh). A thin support film covers the entire grid, typically containing an array of holes with diameters ranging from 1 to 5 µm. Fig. 1a shows a schematic representation of a single mesh of commonly used R 2*/*2 (2 µm diameter holes separating by 2 µm spacing), 200 mesh EM grids.

**Figure 1:**
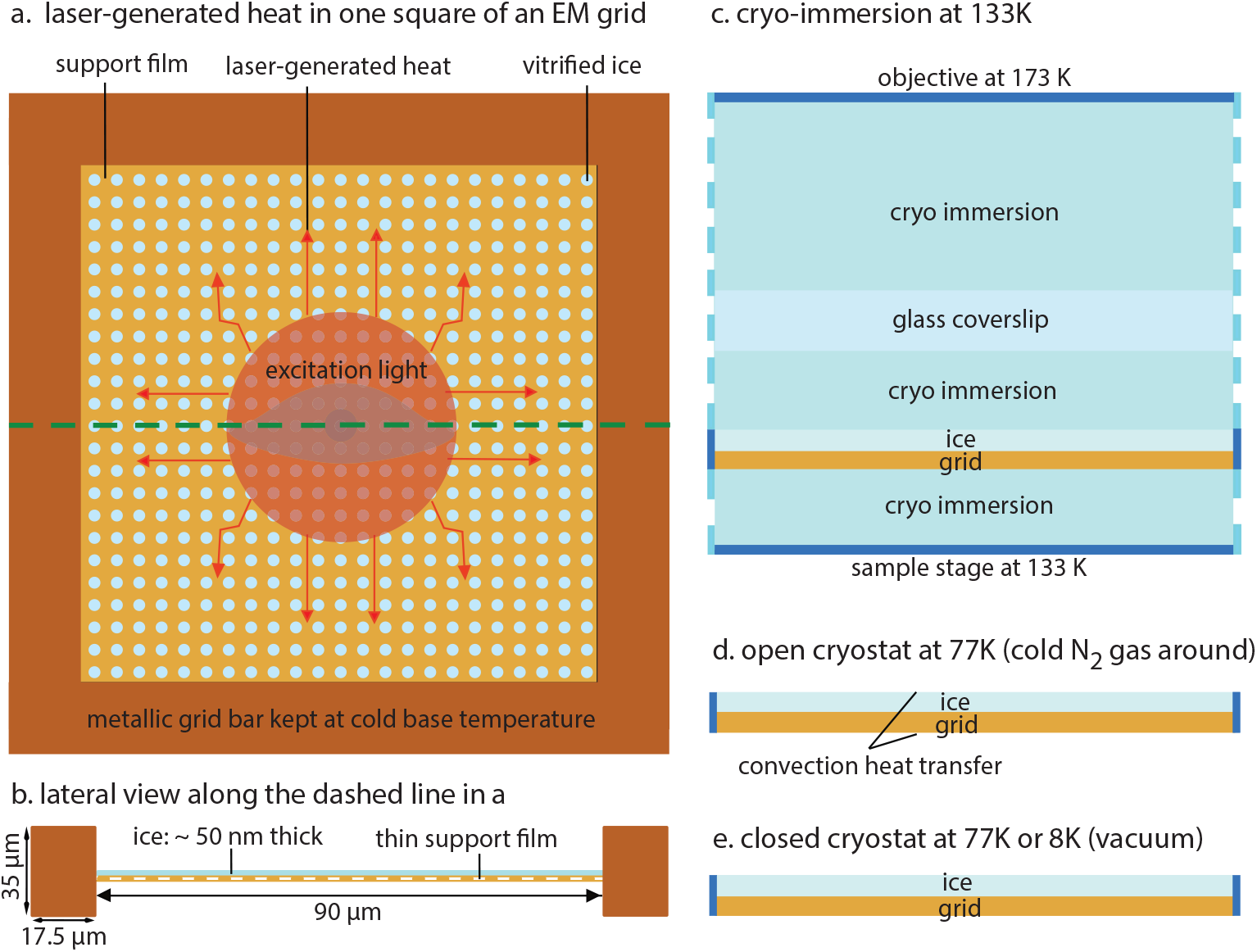
Single mesh of an EM grid and the applied geometries for laser-intensity heating simulations. (a) One square of an R 2/2, 200 mesh TEM grid overlayed with a support film and a frozen biological sample, where the heat generated by absorbed light flows toward the surrounding thick metallic grid bars playing the role of heat sinks. (b) Lateral view of (a) indicating the considerably larger thickness of the grid bar compared to the film and ice. (c) Lateral view of the simulated cryo-immersion geometry (133 K base temperature). The continuous blue line in the geometry illustrates the fixed temperature boundary condition at the upper and lower surface areas, as well as the grid bar. The upper surface, simulated with a temperature of 173 K, represents the temperature of the tip of the cryo-immersion microscope objective. The dashed blue line indicates the surface areas where periodic boundary conditions are assumed. The stage surface positioned 50 µm below the sample is modeled to be at 133 K. (d) Lateral view of sample and film in a LN2-cooled open cryostat with an assumed 77 K temperature at the grid bars (shown as left and right blue lines on the sides) and the normal convection heat transfer provided by the cold N2 gas at the top and bottom of the grid. (e) The same geometry as (d) but without the convection heat transfer, to represent vacuum-based cryostats cooled by LN_2_ or LHe.

Using the finite element method in COMSOL Multiphysics (Materials and methods), we conducted heat transfer simulations to solve the three-dimensional heat equation (1) under thermal steady-state conditions at the level of a single mesh of an R 2*/*2, 200 mesh EM grid:

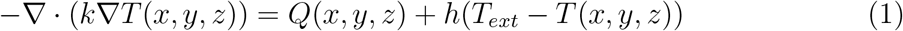

where *k* indicates the heat conductivity of the support film material, ice, and the immersion medium. *T* (*x, y, z*) represents the position-dependent temperature and *Q*(*x, y, z*) is the position-dependent heat generated by light absorption (Materials and methods). *h* denotes the convection heat-transfer coefficient, which we assume to be zero for cryoimmersion and vacuum-based mediums. *Text* denotes the cooling base temperature. As shown in Fig. 1b, the thickness of the metallic grid bars (∼35 µm) is usually over two orders of magnitude greater than the thickness of the support film (™5 100 nm), implying that the grid bars can serve as a heat sink, maintaining a fixed temperature at the base level of the cryostat. In other words, the heat generated by light absorption dissipates toward the grid bars (shown by the red arrows in Fig. 1a). The thickness of the ice layer may vary between different regions of the frozen sample, but we consider it as a flat layer with a thickness of 50 nm on top of the support film considered with a thickness of 12 nm for carbon and 50 nm for gold or silver. The light profile is modeled as a Gaussian shape with full width at half maximum (FWHM) of 35 µm.

We modeled heat transfer for three experimentally available systems: a cryoimmersion cryostat operated at a base temperature of 133 K [31, 32], an open LN2-cooled cryostat at 77 K [12, 13, 15], and a closed (vacuum-based) cryostat at 77 K [28] (Fig. 1c-e).

For the immersion microscope, we assume a geometry as depicted in Fig. 1c. At the top, a surface with a measured temperature of 173 K at thermal steady state (Materials and methods) represents the tip of a cryo-immersion microscope objective that is maintained at room temperature, positioned at 770 µm distance from the EM grid. The lower surface, with a distance of 50 µm from the EM grid and a temperature of 133 K, represents the sample stage. A glass coverslip with a thickness of 170 µm (for the glass-corrected microscope objective) positioned at a distance of 50 µm from the EM grid is included in the geometry to simulate the experimental configuration in the cryoimmersion microscope. Periodic boundary conditions are assumed on the side surfaces of the geometry.

For the open and closed cryostates (77 K or 8 K), we modeled the geometries as shown in Fig. 1d,e. The geometry and boundary conditions modeled for both cryostats are identical, except that we assumed normal convective heat transfer with a coefficient of 20 W/(m K2) at the top and bottom surfaces of the EM grid for the open cryostat, and thermal insulation at these surfaces in vacuum. This accounts for the role of cold N2 gas in heat dissipation [34] (Fig. 1d) as an additional cooling mechanism provided in the open cryostats [26].

Using these geometries, we calculated devitrification thresholds for three grid support materials (carbon, gold, and silver), under laser excitation wavelengths of 488, 561, and 640 nm. Table 1 lists the heat conductivity and light absorption coefficients of amorphous carbon, gold, and silver as gird support film materials, and the heat conductivities of vitrified ice and HFE-7200 at 138 K, whose light absorption were considered negligible. We assume that the metallic material of the mesh has no measurable impact on the laser intensity threshold, as was experimentally verified for copper and gold [27].

**Table 1:** Simulation parameters.

| Material | Heat conductivity at 138 K (W/m·K) | References |
| --- | --- | --- |
| amorphous carbon film | 0.3 | [26, 35, 36] |
| gold film | 72 | [37–39] |
| silver film | 64 | [26, 37, 39] |
| vitrified ice | 1 | [40, 41] |
| glass coverslip | 1 | [33, 42] |
| HFE-7200 | 0.1 (at $\leq 138 \text{ K}$) | [33, 43] |
| liquid propane | 0.2 (at $\leq 138 \text{ K}$) | [44] |
| Material | Optical absorption at 488, 561, and 640 nm (%) | References |
| amorphous carbon | 3.6, 2.8, 1.8 (12 nm thin film) | [45] |
| gold | 50.7, 14, 4.8 (50 nm thin film) | [46, 47] |
| silver | 4.5 (50 nm thin film) | [48] |

According to Equation (1), the laser-generated thermal load is balanced by heat dissipation through the heat conduction of the grid’s support film, the heat conduction of the ice, and the heat conduction or convection from the surrounding medium in non-vacuum cryostats. The excitation intensities that result in a maximum temperature of 138 K are indicated by the simulation readout, and represent the devitrification intensity thresholds.

At 488 nm, the thermal load on carbon is more than ten times lower than on gold due to lower absorption (Table 1). Our simulations show that this relatively low thermal load can be effectively dissipated by the cryo-immersion medium HFE-7200, because of its large cross-sectional area with respect to the support film. Therefore, even at a high base temperature of 133 K of the immersion system, the HFE-7200 medium provides superior heat dissipation for carbon support films (*∼*5.3x) compared to gold films at 488 nm (Fig. 2a). This ratio is reduced by *∼*1.9x at 561 nm, and nearly levels up at 640 nm, in line with the corresponding optical absorption (thermal load) ratios of carbon and gold (Fig. 2a and Table 1).

**Figure 2:**
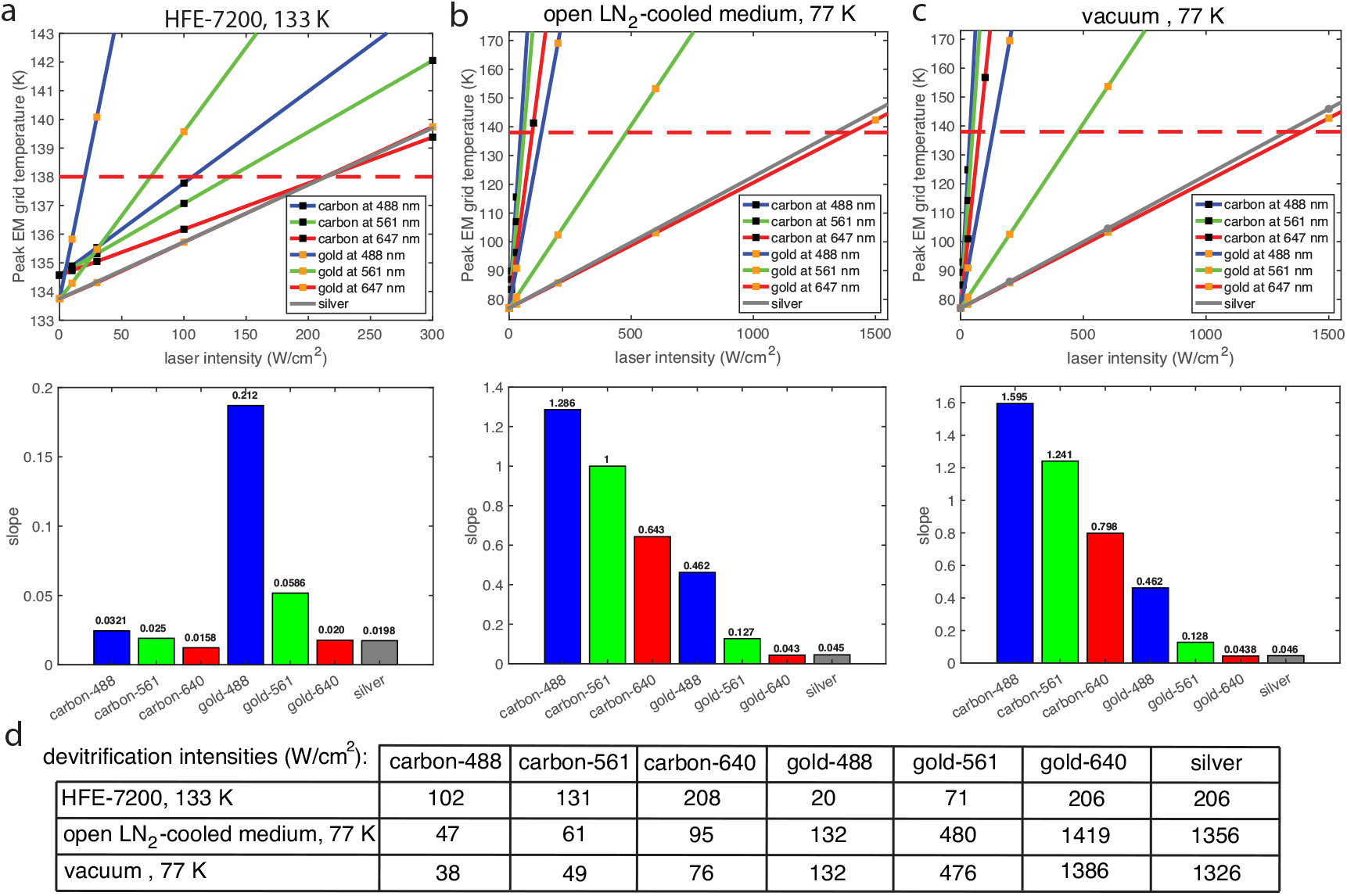
Comparison of maximum safe laser intensities at different immersion mediums, optical wavelengths, and support film materials for a R 2*/*2, 200 mesh EM grid. (a) cryo-immersion at 133 K, (b) LN2-cooled medium at 77 K, and (c) vacuum at 77 K. Lower panels show the slope of the line intensity plots in a-c for a support film material at a certain excitation wavelength in nm (d) the corresponding devitrification laser intensities.

The contribution of convective heat transfer from cold N2 gas is much smaller than the conductive heat transfer by HFE-7200 medium or by the support film, leading to relatively low devitrification laser thresholds for carbon in a cold N2 gas medium (Fig. 2b). Here, the grids with metallic support films tolerate markedly higher laser thresholds compared to those in HFE-7200 medium. For example, almost half of the illuminated light is absorbed by the gold support film at 488 nm, leading to substantial heating.

However, gold also has about 240 times higher thermal conductivity than carbon (Table 1). Furthermore, the heat transfer from the support film to the grid bars is higher at the lower base temperature of LN2 (77 K) compared to the HFE-7200 medium (133 K), where the thermal gradient between the base and devitrification temperatures is only 5 K. Hence, the substantially lower base temperature provided by a LN2-cooled cryo-stage (77 K) allows approximately seven times more efficient heat transfer, effectively managing the larger thermal loads absorbed by gold grids. The gold and silver grids in LN2-cooled open and closed cryostats (77 K) provide up to 6.5 times higher heat transfer (laser thresholds) than the HFE-7200 cryostat at the corresponding wavelengths (Fig. 2). In summary, HFE-7200 enables higher laser thresholds for carbon, while LN2-cooled cryostages allow higher laser thresholds for metallic support films. In addition, the intensity threshold difference between LN2-cooled open and closed cryostats at 77 K is 20% for carbon but only 2% for metallic supports. This suggests that convection heat transfer from cold N2 gas plays a more important role in the dissipation of heat from carbon films. In contrast, for metallic supports, their high thermal conductivity is the primary factor in handling thermal loads (Fig. 2b,c).

To further characterize the effect of the base temperature, we simulated carbon and gold grids at 488 nm in liquid propane, a cryo-immersion medium with a heat conductivity comparable to HFE-7200 but with a base temperature of 86 K [22]. As predicted, the substantially lower base temperature leads to enhanced laser thresholds of 360 W/cm^2^ for gold and 2960 W/cm^2^ for carbon support films.

Unlike silver-coated grids, which are not readily available, gold grid supports have wavelength-dependent, and relatively low, light absorption coefficients at 640 nm compared to 561 nm or 488 nm, thus potentially allowing for higher laser intensity thresholds using 640 nm excitation. However, most of the commonly used fluorophores in SR-cryo-CLEM are excitable with a 488 nm laser [12], with the exception of the red-emitting PAmKate, which can be excited by a 561 nm laser [13, 16]. Despite the lack of a well-characterized fluorophore that can be excited by red (640 nm) lasers for cryoSMLM, we performed simulations at this wavelength because it is particularly relevant for the commercially-available and biocompatible gold-coated grids. Our simulations indeed show that gold grids, when illuminated at 640 nm, permit the highest safe laser intensity across all tested immersion media (cryo-immersion at 133 K, LN2-cooled open, and vacuum cryostats at 77 K), which is attributed to the material’s slightly higher thermal conductivity compared to silver while exhibiting similar light absorption at 640 nm (Fig. 2). This suggests that gold, owing to its biocompatibility and thermal properties, could be an optimal support film, provided suitable (640 nm excitable) fluorophores are developed.

The lower panels in Fig. 2 provide the slope (*α*) of a linear fit to the peak temperature values at different laser intensities across media. This allows calculating the temperature-dependent intensity, I(T), from *I*(*T*) = (*T − T*0)*/α*, where *T*0 denotes the peak EM grid temperature in the absence of heating by light illumination. As expected, *T*0 nearly equals the cooling base temperature in LN2-cooled open or vacuum cryostats where an air objective with a sufficiently long working distance from the EM grid is used. For cryo-immersion in HFE-7200, *T*0 is slightly higher than the stage base temperature due to the axial temperature gradient from the grid surface to the objective tip (133.9 K for gold and silver, and 134.7 K for carbon). Fig. 2d lists the corresponding devitrification intensities.

To directly compare our simulations with experimental results in the literature, we recalculated the laser intensity thresholds using the geometries and parameters in previous studies. In an open cryostat system, for R 1.2*/*1.3, 300 mesh gold support at 488 nm, R 2*/*2, 200 mesh gold support at 561 nm, and C-Flat (R 2*/*1-2C, 200 mesh) carbon support at 488 nm, the recalculated thresholds were 190 W/cm2, 480 W/cm2, and 48 W/cm2. These values agree well with the experimental numbers reported of 186 W/cm^2^ [27], 250-300 W/cm^2^ [26], and 28.5 W/cm^2^ [12], respectively.

Next, we experimentally tested devitrification of EM grids under conditions that match our simulation. For these experiments, we used 200 mesh size grids with R 2*/*2 gold support films or with R 2*/*1 carbon or silver films. Our experiments, designed to validate some of the key simulation results, included. (1) a cryo-immersion microscope operated at a base temperature of 133K for laser dosing of grids with carbon and gold support films illuminated at 488 nm. (2) a LN2-cooled open cryostate custom-built microscope (Materials and methods) operated at 95K for laser dosing of gold-support grids at 640 nm. (3) a vacuum-based microscope [28] operated at 77K for laser dosing of silver-coated grids. Figure 3a shows a table with the simulated laser intensity thresholds for the relevant experimental specifications, i.e., the support film material and dimensions, immersion medium, base temperature and laser wavelength.

**Figure 3:**
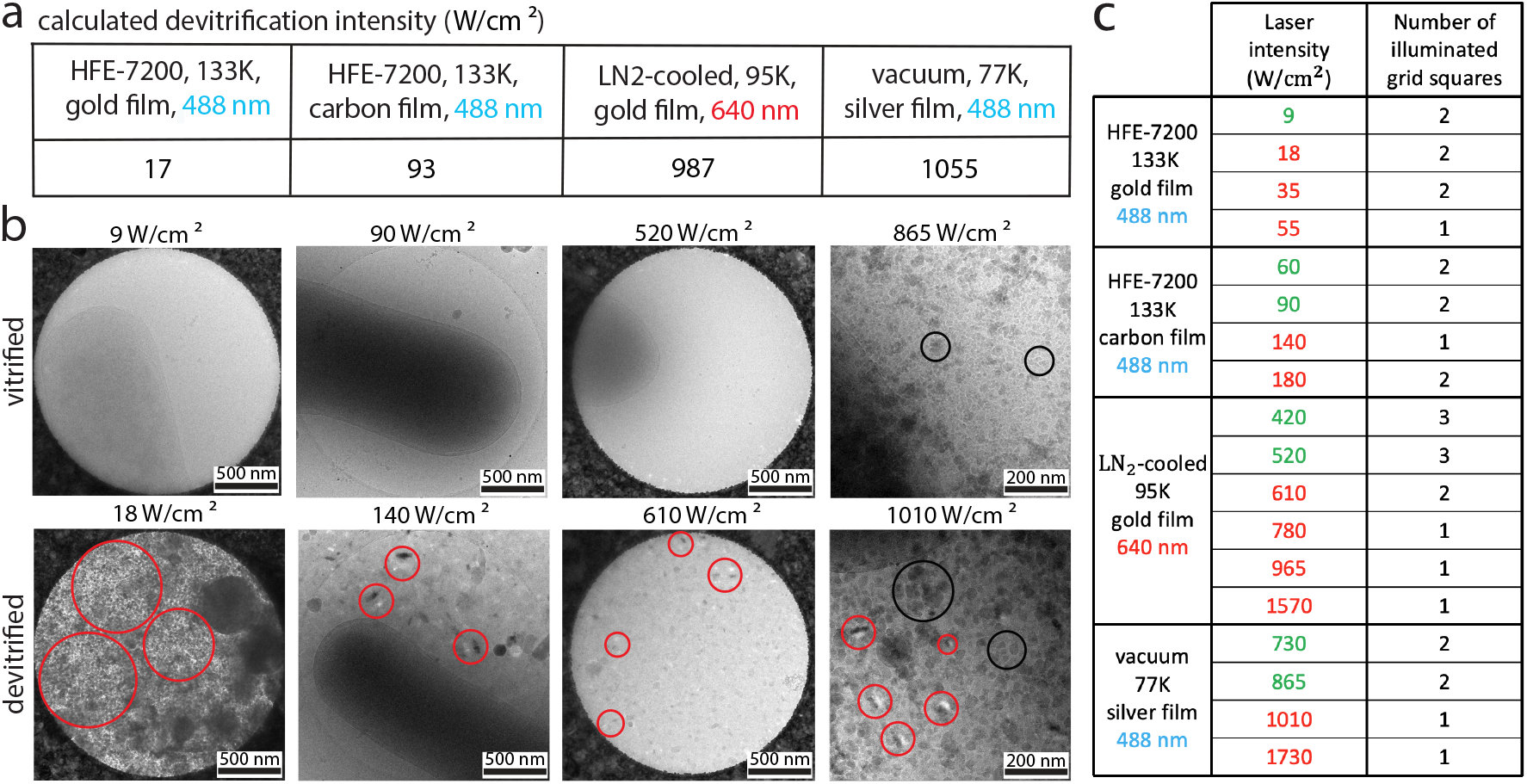
Laser intensity thresholds for ice devitrification in HFE-7200, LN2-cooled open and closed mediums. a) Simulated laser intensity thresholds for devitrification on grids with carbon and gold support films illuminated with a 488 nm laser in HFE-7200, on a grid with a gold support film illuminated with a 640 nm laser in an LN2-cooled cryostat, and on a silver support film illuminated with a 488 nm in a vacuum cryostat. b) Cryo-EM micrographs acquired after laser illumination corresponding to the condistions described in (a). The top and bottom rows display vitrified and devitrified ice, respectively, around the specified peak laser intensities. The shown grid holes are selected from the centers of grid squares where the peak intensity of the Gaussian laser beam was applied. c) Laser intensities applied to each grid with corresponding statistical data. The laser intensities colored in green and red represent conditions resulting in vitrified and devitrified ice, respectively.

To validate the simulation results, we vitrified *E. coli* cells on the grids (Materials and methods) and imaged them with cryo-EM to confirm initial ice vitrification, and to map the positions of the grid squares with thin samples. Subsequently, the same grids were loaded into the different cryostats. We illuminated different grid squares with laser intensities slightly below and above the predicted thresholds. The ice status at these positions was evaluated again using cryo-EM. Fig. 3b shows cryo-EM micrographs corresponding to the experimental conditions in Fig. 3a after approximately 30 seconds of laser exposure at the intensities indicated above each image. The top panels display the highest laser intensity at which the ice remained vitrified, corresponding to the safe operating limit. The bottom panels illustrate where devitrification occurred (marked with red circles).

For the cryo-immersion cryostat in HFE-7200, we observed a laser threshold below 18 W/cm^2^ for grids with gold supports at 480 nm. In contrast, for grids with carbon supports in the same cryostat, the threshold ranged between 90 and 140 W/cm2, showing good agreement with our simulation results.

For the open cryostat and grids with gold support films, experimental validation shows that devitrification occurs between 520 and 610 W/cm^2^ at 95K when using a 640 nm laser (Fig. 3b). For context, previous studies [27], [26] reported devitrification thresholds for gold supports of around 250 W/cm^2^ at 488 nm (R1.2*/*1.3, 300 mesh) and 250 *−* 300 W/cm^2^ at 561 nm (R2*/*2, 200 mesh), both measured at 77 K. Adjusting for our base temperature of 95 K for R2*/*2 200 mesh grid, these thresholds are expected to decrease to roughly 125 W/cm^2^ (488 nm) and 210 *−* 250 W/cm^2^ (561 nm). Thus, the devitrification threshold at 640 nm is approximately three times higher than at 561 nm and approximately five times higher than at 488 nm, demonstrating the wavelength-dependent nature of devitrification on gold support film grids.

Our experiments in the vacuum-based cryostat show that vitrification is maintained even at relatively high laser intensities using grids with custom-coated silver thin films (Fig. 3b). A previous study demonstrated the threshold intensity of 1 *−* 1.25 kW/cm^2^ for silver thin films in an open LN2-cooled cryostat at 77 K [26]. Overall, the intensity thresholds in vacuum are slightly lower than those observed in open cryostats at the same base temperatures, presumably due to the lack of the additional minor contribution of cold N2 gas to heat dissipation. The black circles in Fig. 3b show ice contamination that occurred during the loading and unloading of grids during the experiments using the cryo-vacuum cryostat. Despite this, the appearance of crystallized (devitrified) ice (red circles) remains distinct from the contamination. Fig. 3c summarizes the laser intensities applied in each experiment, where green indicates safe illumination (vitrified ice) and red indicates devitrified or sublimated ice.

Overall, our experimental results align with the simulation predictions. The discrepancies observed for the gold-support grid at 640 nm (95 K, open cryostat) likely result from slightly higher temperatures at the grid position, located 7 mm above the position of the temperature sensor inside the sample holder. They may also arise from minor variations in light absorption and thermal conductivity of the support film used in the experiments.

Having characterized the key role of the cryostat base temperature in laser-induced ice devitrification, we next explored the scenario of a vacuum-based, helium-cooled cryostat that provides a base temperature of 8 K [28, 29]. As shown by Hoffmann et al. [28], fluorophores not only produce higher signal-to-background ratios at 8 K compared to 77 K, but also often exhibit a longer off/on time ratio (the so-called dynamic contrast ratio). This facilitates more efficient single-molecule detection and cryo-SMLM of densely-labeled samples with higher accuracy and localization precision. In thin films, the mean free path of heat carriers (such as phonons or electrons) is limited by the finite grain size of the material. This increased boundary scattering leads to lower thermal conductivity compared to the bulk material, where heat carriers experience fewer such constraints [49, 50]. At ultra-low temperatures, we must account for the temperature dependence of the heat conductivity, *k*(*T*). The heat conductivity in the temperature range of 77-138 K can be approximated as nearly constant [26, 27]. However, to explore the impact of a possible temperature dependence, we also considered a scenario where *k*(*T*) follows a linear decrease in this range. The temperature-dependent conductivity of thin films at ultra-low temperatures can be approximated using the Wiedemann-Franz law, by applying the measured temperature-dependent electrical resistivities [51]. Based on this, we modeled a linear drop in heat conductivity from 138 K to 77 K and from 138 K to 8 K using *k*(*T*) = *cT* where *c ≈* 0.46 W/m*·*K2, and *k*(*T* = 138 K) = 64 W/m·K for a silver film (Table 1). In Fig. 4a, we compare the calculated laser intensity thresholds for silver support in vacuum-based cryostats at 77 K and 8 K under both constant and temperature-dependent thermal conductivity assumptions. At 77 K, assuming a temperature-dependent conductivity results in a threshold about 24% lower (1.065 kW/cm2, shown in orange) compared to the constant conductivity case (1.325 kW/cm2, shown in red). This difference becomes more pronounced at 8 K, where the threshold is reduced by 86%. Despite the reduced heat conductivity of thin films at lower temperatures, the laser intensity threshold at 8 K (1.52 kW/cm2) remains 43% higher than at 77 K (1.065 kW/cm2), primarily due to the larger (130 K) temperature gradient to the devitrification threshold. Fig. 4b presents the laser intensity thresholds for carbon, gold, and silver grid supports at various wavelengths, calculated with temperature-dependent heat conductivity at an 8 K base temperature. These findings highlight the impact of temperature-dependent thermal conductivity at ultra-low temperatures on laser intensity thresholds, showing that while operating at 8 K offers some advantages for cryo-SMLM and single-molecule detection, the intensity threshold improvement over 77 K is not highly significant.

**Figure 4:**
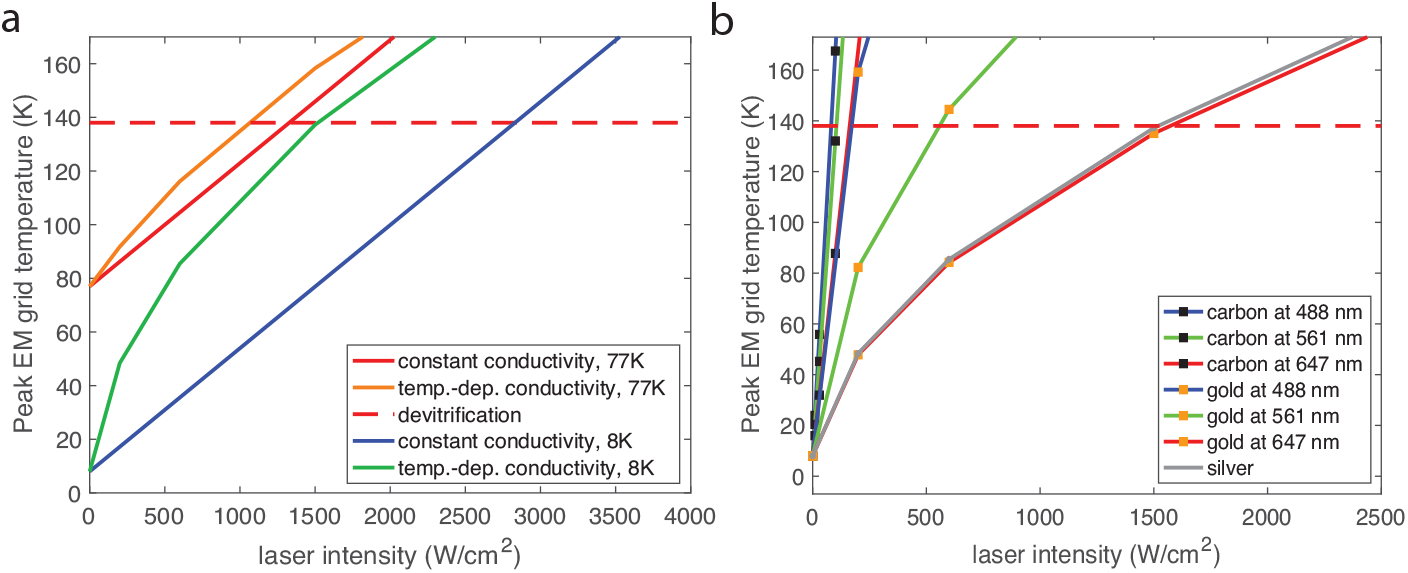
Comparison of light-generated heat transfer with constant and temperature-dependent heat conductivity on a vitrified EM grid for a vacuum-based cryostat (a) at 77 K and 8 K using a silver-coated grid and (b) at 8 K for different support film materials with temperature-dependent heat conductivity.

In summary, our study shows that cryo-immersion media improve heat dissipation for optically low-absorbing support films, while the high thermal conductivity of metallic support films allows them to withstand higher laser intensities in cryostats with low base temperatures. Additionally, metallic supports continue to efficiently dissipate heat in vacuum-based cryostats, making them ideal for a variety of cryo-SMLM setups.

## 3. Discussion

In this work, we investigated critical factors that affect devitrification thresholds in SR-cryo-CLEM. Through finite element simulations, we systematically evaluated the roles of base temperature, immersion medium, and the thermal and optical properties of grid support film materials, and experimentally validated key findings in three different cryoSMLM setups.

We show that cryo-immersion media allow for higher light intensities for low-absorbing EM support films like carbon, compared to LN2-cooled cryostats operating at 77 K or 8 K. This finding could be valuable for applications where the carbon-support films, often implemented together with silicon dioxide for increased rigidity in preparations of cellular samples, are sometimes preferred over delicate gold films. Another potential benefit of a cryo-immersion medium could include reduced contamination during the long acquisitions required for cryo-SMLM, compared to open LN2-cooled cryostats [32]. Despite the higher photon collection efficiency (NA=1.15) and higher heat conductivity to the surrounding medium provided by cryo-immersion microscopes, for metallic-support grids, current cryo-immersion implementations with near-devitrification base temperatures only permit about 15% of the light intensities achievable with LN2-cooled cryostats at 77 K base temperature. In other words, highly conductive metallic support films, such as gold, benefit significantly from the lower base temperatures in LN2-cooled or LHe-cooled stages, allowing for safer operation at higher laser intensities.

Our study further validates the role of support film material in managing the tolerable laser dose. Metallic support films, especially gold, with their high thermal conductivity and low light absorptions at 640 nm, enable safe laser intensities that are higher than those of commonly used carbon supports in cellular cryo-EM. These properties position gold as an optimal biocompatible support material, underscoring the need for future development of far-red emitting fluorophores. Furthermore, our findings indicate that vacuum-based cryostats operating at 77 K enable the use of laser doses comparable to those used in open cryostats at the same temperature, particularly when metallic-coated grids are utilized. Additionally, vacuum cryostats offer a key advantage in cryo-SMLM by significantly reducing ice contamination, which can accumulate over long cryo-SMLM data acquisition times and negatively affect image quality.

The heat transfer simulation program developed in this study opens a path for investigating a broad range of possible practical applications. This program provides a tool for precise modeling of thermal dynamics in cryo-SMLM, enabling researchers to optimize experimental designs for available microscope setups and specific imaging conditions. By incorporating parameters such as base temperature, immersion medium properties, EM grid support film material and geometry, ice thickness, and illumination laser properties, this tool can guide experimentalists in designing cryo-SMLM investigations that maximize safety and performance.

Finally, this study draws attention to the practical challenges and design considerations for cryo-SMLM systems. Vacuum-based cryostats at 8 K, while offering the lowest base temperatures, require careful sample stage design to ensure efficient heat dissipation via conduction. Open LN2-cooled systems, while simpler and more cost-effective, face challenges such as vibrations due to LN2 boil-off, which can compromise mechanical stability in the relatively long acquisition times in cryo-SMLM. Cryo-immersion methods, while providing high heat dissipation, are limited by their base temperatures due to the freezing point of cryo-immersion media.

In conclusion, this work provides quantitative insights into the factors governing devitrification in cryo-SMLM, emphasizing the importance of carefully considering combinations of base temperature, immersion medium, and support film material properties. Our findings not only advance the understanding of thermal dynamics in cryo-CLEM, but also offer practical tools and guidelines for designing optimized experimental setups.

## 4. Materials and methods

### 4.1 Details of heat transfer simulations

We conducted heat transfer simulations at a thermal steady state using COMSOL Multiphysics version 6.0. The spatially distributed heat source generated by an incident laser with a Gaussian profile was modeled as 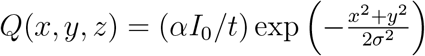 where *α*, 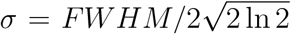 correspond to the optical absorption coefficient of the support film material, the thickness of the support film, the peak intensity at the center of the beam and the standard deviation of the Gaussian profile, respectively. The finite element method simulations employed a mesh based on free tetrahedral physics with a normal element size. This meshing strategy was chosen to ensure accurate resolution of sharp thermal gradients near the laser focus while maintaining computational efficiency. To quantify the risk of devitrification, the primary output of the simulation was the maximum temperature reached within the grid and the volume of the ice. This readout provided a direct metric for comparing thermal behaviors across support film materials, ice, and immersion media. The COMSOL programs provided for vacuum or LN2-cooled media (file: N2-77K.mph), HFE-7200 media (file: HFE-7200-133K.mph), and detailed instruction on how to use them are available in (Github, ries-lab/devitrificationSim).

### 4.2 Sample preparation and cryo-EM imaging

Quantifoil SiO2/carbon-coated copper 2*/*1 200 mesh grids and UltrAuFoil all-gold 2*/*2 200 mesh grids (Quantifoil Micro Tools) were plasma cleaned with a Fischione M1070 (90:10 Ar:O2 gas mix) for 1 minute before sample application. Alternatively, silver-coated grids were fabricated using a similar procedure to Dahlberg et al. [26]: Quantifoil copper 2*/*1 200 mesh grids (Quantifoil Micro Tools) were harshly plasma cleaned, 5 nm of nickel was deposited on grids as a wetting layer, and 50 nm of silver was subsequently deposited with a Leica ACE600 (Leica Microsystems).

*E. coli* DH5a cells were grown to log phase in lysogeny broth (LB) medium, and 3 microliters of sample were applied to EM grids. Samples were blotted from the back for 2 seconds and subsequently plunge frozen using liquid ethane (88 K) on a Leica GP2 plunger at 40% humidity.

Cryo-EM micrographs were collected using SerialEM [52], and were acquired on a Talos Arctica 200 keV microscope (Thermo Fischer Scientific) equipped with a Falcon 3 detector (Thermo Fischer Scientific) at 8.441 Å/pixel, a 100 µm objective aperture inserted, approximate fluence 10 e^*−*^*/*Å2, and 10 µm nominal defocus. Alternatively, for analysis of the devitrification threshold in the vacuum cryostat, micrographs were acquired on a Titan Krios microscope at 300 keV (Thermo Fischer Scientific) equipped with a Gatan K3 detector and BioQuantum post-column energy filter (Gatan), at 8.858 Å/pixel, with a 100 µm objective aperture inserted, approximate fluence 10 e^*−*^*/*Å2, and with 2-3 µm nominal defocus.

### 4.3 Front lens temperature measurements for a cryo-immersion objective

As a boundary condition for the simulation of the cryo-immersion medium, we measured the temperature of the front lens of the cryo-immersion objective (63x, 1.15 NA), as described in [32]. In the experimental setup, a PT1000 temperature sensor (dimensions: 2.0 × 2.0 × 1.3 mm) is glued to a cover glass and placed in the optical path at the sample position and embedded in a droplet of HFE-7200. The objective body is heated to room temperature with the heater as described by Faoro et al. [31] and Faul et al. [32], while the tip is immersed in the cold HFE-7200, cooled by the sample holder (133 K). The front lens was then positioned 150 µm above the sensor to avoid mechanical contact (working distance 600 µm). However, due to the sensor height of 1.3 mm, the total distance to the cold sample was increased by 1 mm compared to standard imaging conditions. At equilibrium, the measured temperature near the front lens was 177 K. Due to the increased distance from the lens to the sample, the temperature tends to be overestimated and the actual temperature is lower. We therefore assume 173 K as a boundary condition for the simulation.

### 4.4 Laser dosing experiments using a cryo-immersion microscope operating at 133K

Cryo-immersion light microscopy was performed as previously described [31, 32]. The cryo-immersion objective (Zeiss LD C-Apochromat, 63x, 1.15 NA) was mounted on an upright microscope (Axio Scope.A1, Zeiss), equipped with a custom-built cryo-stage for cryo-immersion microscopy. HFE-7200 was used as the cryo-immersion medium. Light microscopy images were captured using an sCMOS camera (Panda 4.2, PCO).

For brightfield and epifluorescence, the SpectraX lamp (Lumencor) was used in combination with the quad-band filter set (dichroic ZT405*/*488*/*561*/*640rpcv2-UF2 and emission filter ZET405*/*488*/*561*/*640mw2, AHF). The 488 nm laser (C-FLEX laser combiner, Hübner Photonics) is incorporated via a custom-made beam splitter adapter mounted on the microscope field diaphragm (BS037 50:50, Thorlabs). The laser is guided through the fiber (kineFlex, Qioptiq), a 3x beam expander (GBE03-A, Thorlabs), and an aperture to adjust the spot size, and focused into the back focal plane of the objective. The laser intensity was calibrated by measuring the power at the back focal plane using a power meter (PM100D, Thorlabs) and a photodiode (S121C, Thorlabs). To compensate for the objective, a transmission loss of 20% was taken into account according to the data sheet of the objective. The spot size was adjusted to *∼*40 µm FWHM. Prior to imaging, samples were embedded in HFE-7200 and transferred to the microscope. To ensure proper cooling of the samples mounted in autogrid cassettes, the same shuttle design, as described in [31, 32], was used. After imaging, the sample was removed from the shuttle, and HFE-7200 was removed by dissolution in liquid ethane at −138°C [32].

### 4.5 Laser dosing experiments using a microscope with an open cryostat operating at 95K

We used a custom-built cryo-fluorescence microscope to navigate, image, and illuminate grid squares. A 640 nm laser (LuxX, Omicron, 140 mW) coupled to a multimode optical fiber (150×150 µm, 0.39 NA, M1034L05) provided illumination. To achieve uniform illumination, we reduced beam speckling by mechanically shaking the fiber with a vibration motor (DC 1.5-6V, 16500 rpm, Sourcingmap) [53]. The fiber output was collimated with an aspheric lens (f = 10 mm) and focused using a doublet lens (f = 200 mm) at the back focal plane of an upright objective (CFI TU Plan Apo EPI 100×, 2 mm WD, Nikon). The field of view size was tuned to 35 µm FWHM. Fluorescence signals collected by the objective passed through a quad-band dichroic mirror (F73-410, AHF) and were focused by a tube lens (f = 200 mm, TTL200-A, Thorlabs). A 4f relay system (two lenses: f = 200 mm, f = 100 mm) projected the image onto an sCMOS camera (ORCA-Fusion BT, Hamamatsu).

The cryostat consists of a cage structure with Polyether Ether Ketone (PEEK) side and top plates and a copper base plate, which secures the translation stages and sample holder. The cage is mounted on an aluminum plate with a 100×100 mm opening and fixed to an optical breadboard (600×600 mm, Thorlabs). The objective, attached via a copper adapter to the PEEK plate, is heated with foil heaters and monitored by a temperature sensor to maintain 297.3 K, preventing frost accumulation. A plastic cap thermally insulates the front of the objective. The entire cage is enclosed in a heat-insulating container, into which liquid nitrogen (LN2) is pumped via a micro-dosing system (Norhof, model 915). LN2 is maintained just below the base plate to ensure efficient heat transfer to the translation stages and sample holder. The system includes two lateral translation cryo-stages (ANPx311, Attocube) and a vertical lifting cryo z-stage (ANPz-102, Attocube), all operable at cryogenic temperatures.

The sample holder temperature is regulated at 95 K using a controller module (model 335, LakeShore Cryotronics) and a heat-conducting device (ATC100/70, Attocube), consisting of two copper plates connected by copper braids, a heater, and a temperature sensor. The heat-conducting plate with the sensor and heater is mounted on the z-stage, while the second plate is fixed to the base plate for enhanced thermal conductivity. LN2 pumping is optimized to minimize stage temperature oscillations (±0.03 K) during dosing. We used Leica copper cartridges (Leica Microsystems) as sample holders, each containing two slots with fine copper springs to secure EM grids mounted in autogrid cassettes. A magnet at the bottom of the cartridge ensures firm contact with the copper adapter plate mounted on the heat-conducting plate. EM grids were loaded and unloaded using a commercial cryogenic transfer shuttle (Leica Microsystems).

### 4.6 Laser dosing experiments using a microscope with a vacuum-based cryostat operating at 77K

We conducted the laser dosing experiments in a vacuum-based cryogenic wide-field microscope detailed in [28]. We obtained the optical transmission factor of the incorporated microscope objective (Nikon CFI L Plan EPI CRB 100x NA=0.85 air objective) for a 488 nm laser (4 W, Coherent, Genesis CX STM 488nm 4W) by measuring the laser power before and after the microscope objective, giving the transmission factor of *∼* 0.845. The measured laser powers just behind the microscope objective (at the back focal plane) were multiplied by the transmission factor to estimate the delivered laser power at the sample plane inside the vacuum cryostat. The size of the field of view was adjusted to *∼* 35 µm FWHM. The cryostat of the cryogenic fluorescence microscope kept at 77 K, a vacuum of *∼* 2 *×* 10*−*5 Torr. EM grids were loaded and unloaded using a commercial cryogenic vacuum transfer system (Quorum Technologies, PP3010T).

## Acknowledgements

This work was supported by the Chan Zuckerberg Initiative (CZI) Visual Proteomics Program (Grant No. 2021 *−* 234620), the EMBL, and the European Research Council under the European Union’s Horizon 2020 Research and Innovation Program (Grant No. 772441). The authors acknowledge the facilities provided by the EMBL Cryo-EM Platform and the EMBL Imaging Centre, the support of Harald Hess, Gleb Shtengel, and James Seyforth at the Janelia Research Campus for their guidance and assistance in conducting the dosing experiments with the vacuum cryostat microscope, and the cryoelectron microscopy facility team at Janelia, especially Zhiheng Yu, Momoko Shiozaki, and Xiaowei Zhao, for their help during imaging. Finally, we thank the group members of J.M., J.R., and T.P.B. for their insightful discussions and valuable input throughout this work.

## Declaraion of conflict of interest

J.M. is on the editorial board of JSB. All authors declare no conflict of interest.

